# GDF15 knockout does not substantially impact perinatal body weight or neonatal outcomes in mice

**DOI:** 10.1101/2024.04.30.591359

**Authors:** Molly C. Mulcahy, Noura El Habbal, JeAnna R. Redd, Haijing Sun, Brigid E. Gregg, Dave Bridges

## Abstract

Growth differentiation factor-15 (GDF15) increases in circulation during pregnancy and has been implicated in food intake, weight loss, complications of pregnancy, and metabolic illness. We used a *Gdf15* knockout mouse model (*Gdf15^-/-^* ^)^ to assess the role of GDF15 in body weight regulation and food intake during pregnancy. We found that *Gdf15^-/-^* dams consumed a similar amount of food and gained comparable weight during the course of pregnancy compared to *Gdf15^+/+^* dams. Insulin sensitivity on gestational day 16.5 was also similar between genotypes. In the postnatal period, litter size, and survival rates were similar between genotypes. There was a modest reduction in birth weight of *Gdf15^-/-^* pups, but this difference was no longer evident postnatal day 3.5 to 14.5. We observed no detectable differences in milk volume production or milk fat percentage. These data suggest that GDF15 is dispensable for changes in food intake, and body weight as well as insulin sensitivity during pregnancy in a mouse model.

## Introduction

Growth-like differentiation factor-15 (GDF15), a Transforming Growth Factor-ß superfamily member, placental derived growth factor, and cytokine, was discovered in 1997 and dubbed macrophage-inhibiting cytokine-1 (MIC-1) (1). Circulating levels of GDF15 in adults vary based on sex, age, disease status, and physiological state. A large sample study from Scotland found that levels of circulating GDF15 increase with age in both men and women and tended to be higher in those who had cardiovascular disease, cancer, or diabetes (2). GDF15 increases in response to many stressors including cardiac injury (3), cachexia of cancer (4), mitochondrial stress (5), intense exercise (6), and most relevant to this work, during pregnancy (2,7–10).

Preclinical work with knockout or knockdown models has highlighted the role of GDF15 in body weight regulation (11), appetite (12), and emesis (13). In rodents, the effect of GDF15 antagonism through antibodies or knockout on food intake depends on diet. When consuming a high fat, high sucrose diet, food intake and body weight increases (12,14); however, when consuming a chow diet, food intake remains similar to wildtype animals (14). These models show that GDF15 acts through the GFRAL receptor found in the area postrema of the brain. The role of GFRAL in body weight and food intake has been just as critical as GDF15. There is evidence of a positive association between GFRAL positive neurons and fat mass/body weight gain (12). Interrupting GFRAL receptors in preclinical models does not produce consistent results on weight and feeding behavior. One model showed ablating GFRAL in mice resulted in smaller mice at the beginning of the study that then developed increased food intake and weight gain from eating a hyperpalatable diet (15). Another noted no differences in food intake, weight accretion, or in size at the onset of the experiment (16).

Overexpression or pharmacologic administration of GDF15 induces weight loss through reductions in food intake (11,15,16). GDF15 elevation also results in nausea-like behavior in mice and emesis in shrews (13), reduced changes in food preferences (17), or a decrease in meal size (18). As such, evaluating *Gdf15* for its capacity to abrogate dysmetabolism is currently being explored.

During human pregnancy, GDF15 increases across gestation and reaches its highest levels during the third trimester (7,8,10,19). *Gdf15* is heavily expressed in placental trophoblasts, is secreted into parental circulation, and is present in amniotic fluid (19). In fact, the majority of GDF15 in circulation is derived from the fetus (20). Despite these pregnancy-related increases, details on the function of GDF15 in pregnancy are just emerging. GDF15 has been linked to several complications and conditions that can arise during pregnancy. Lower levels of GDF15 during early pregnancy have been found in patients who later suffered miscarriage (21). GDF15 levels have also been linked to gestational weight gain, with elevated levels negatively associated with cumulative gestational weight gain (22). Petry and colleagues found pre-pregnancy BMI was inversely related to GDF15 levels during pregnancy (23). Differing levels of GDF15 have been detected in concert with complications of pregnancy. In several cases, the epidemiological data is conflicting. For example, pre-eclampsia, a life-threatening complication involving critically high blood pressure and protein loss in urine, has been found to be associated with reductions (8), increases (10,24), and no changes (9) in GDF15 in serum compared to non-preeclamptic, normotensive parents. Similarly, some studies found that GDF15 is higher in pregnancies complicated by gestational diabetes (GDM) (25), or type 2 diabetes (T2DM) (10), while others found it is only significantly increased in pregnancies that are complicated by type 1 diabetes (T1DM) but not T2DM or GDM (26). GWAS have indicated that *GDF15* variants in humans are associated with hyperemesis gravidarum (HG), an extreme form of nausea and vomiting of pregnancy (27,28). Recent evidence supports high levels of GDF15 in circulation pre-pregnancy may be protective for HG and further elucidates the causal role of GDF15 in severe nausea and vomiting of pregnancy (20). Given the sometimes-conflicting human data, we sought to define the effects of *Gdf15* loss of function during the course of murine pregnancy, including effects on weight gain, food intake, insulin sensitivity, and neonatal outcomes.

## Materials and Methods

### Animal Husbandry

Animals from both studies described below were housed in a temperature and humidity-controlled facility with a 12-hour light: dark cycle, with lights on being zeitgeber time (ZT) 0 and lights off being ZT 12. All protocols were approved by the Institutional Animal Care and Use Committee of the University of Michigan.

### Insulin resistance of pregnancy study

Virgin female C57BL/6J (RRID: IMSR_JAX:000664) mice were purchased from The Jackson Laboratories. Mice were allowed to acclimatize for two weeks to the temperature and humidity-controlled facility with *ad libitum* access to water and laboratory chow diet (CD, Picolab Laboratory Rodent diet 5L0D; 5% of Calories from fat, 24% from protein, 71% from carbohydrates). After acclimatizing, females were randomized into three groups, non-pregnant females (n=7), pregnant females (n=7), and pregnant females exposed to dexamethasone (1mg/kg/day Sigma-Aldrich catalog #D2915-100MG) in drinking water (n=7). One week after experimental treatment began, males were introduced to the dam’s cage and allowed to remain until gestational day 19.5. Body weight and food intake measurements occurred weekly from randomization until birth, or until age-matched dam delivered (**Figure 1A**).

**Figure 1:**
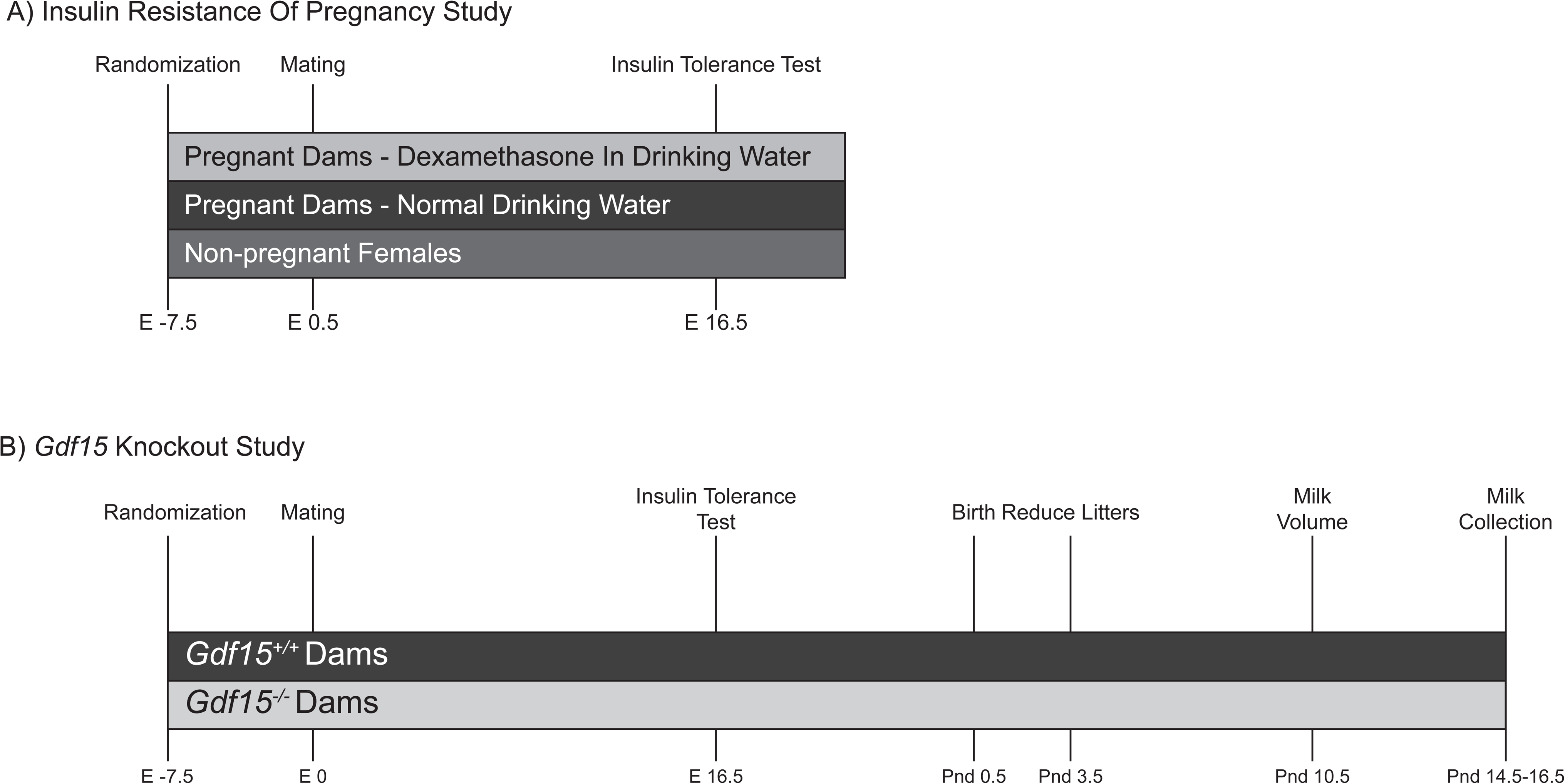
Schematic of Experimental Manipulations. A) Insulin resistance of pregnancy study, comparing age-matched females in 3 groups; non-pregnant females (n=7), pregnant females given plain drinking water (n=7), pregnant females given 1.0 mg/kg dexamethasone in drinking water (n=7). B) *Gdf15* Knockout study in pregnancy. *Gdf15^+/+^*females (n=6) were mated with *Gdf15^+/+^* males. *Gdf15^-/-^*females (n=7) were mated with *Gdf15^-/-^* males. Food intake and body weight was measured weekly from one week before mating until 14-16 days after pups were born.

### GDF15 study

Male and female *Gdf15* null animals (*Gdf15^-/-^*) were previously described (17). Null animals were generated using CRISPR Cas-9 deletion of Exon 2 of *Gdf15*. Exon 2 (translational start site), which we ablated, is present in every known *Gdf15* transcript. We chose to study *Gdf15* wildtype (*Gdf15^+/+^*) mated pairs compared to *Gdf15^-/-^*pairs because comparing littermates of *Gdf15^+/-^* pairs would result in potential placental contributions to GDF1*5* in dam serum as the feto-placental unit provides a substantial amount of GDF15. To limit genetic drift all homozygous parents were direct offspring of heterozygous crosses. We combined homozygous pairs, resulting in homozygous genotype progeny and placentae. Adult virgin female mice (*Gdf15*^-/-^n=8, *Gdf15*^+/+^n=6), between 45 and 119 days old (mean 82 days), were singly housed with *ad libitum* access to water and a CD. Weekly food intake and body weight measurements began after single housing dams and continued throughout the experiment (**Figure 1B**). After one week of food and body weight monitoring, males of like-genotype were introduced into the dam’s cage. Males were allowed to remain in the breeding cage until a copulatory plug was identified, indicating pregnancy (E0.5). Body weight and food intake measurements continued weekly through gestation to postnatal day (PND) 14.5. Their resultant offspring and their placentae were homozygous *Gdf15^+/+^*and *Gdf15^-/-^* and were studied until PND 14.5.

### Genotyping

At 14 days of age, a tail clip was collected and digested in 100uL of PBND lysis buffer (10 mM Tris pH 8.0, 150 mM NaCl, 10 mM EDTA, 0.1% SDS and 1 mg/ml proteinase K) at 55°C overnight, then 85°C for one hour. Digested DNA samples were amplified with DreamTaq Green Master Mix (ThermoFisher Scientific, Catalog #K1081) to generate PCR product. Genotyping by PCR was conducted with 2 forward and one reverse primer sets (forward 1: 5’ GAT TCC CGC CCG AAT TAG C 3’, forward 2: 5’ CCG AAT TAG CCT GGT CAC CC 3’, Reverse: 5’ ATC CGT CCT ACT CTG GCT AAG 3’). Initiation of PCR was at 95 °C for 3 minutes, followed by 38 cycles of denaturation (95°C for 30 seconds), annealing (60°C for 40 seconds), and elongation (72°C for 1 minute), and a final amplification step at 72°C for 5 minutes. PCR product resulted in 2 visible bands, one at 200bp *Gdf15^-/-^* and another at 600bp *Gdf15^+/+^.* Mice with both bands were considered *Gdf15^+/-^.* Dam genotype was secondarily confirmed via maternal serum ELISA (**Supplemental Figure 1**).

### Insulin tolerance tests

On E16.5, dams underwent intraperitoneal (IP) insulin tolerance testing (ITT) (29). Dams were placed in clean cages without access to food but with *ad libitum* access to water at ZT 2. Dams were fasted for 6 hours (ZT2-ZT8). Baseline blood glucose was assessed using a tail clip and a handheld glucometer (OneTouch Ultra). After initial blood glucose measurement, an IP injection of insulin was administered (Humulin, u-100; 0.75U/kg lean mass). Blood glucose was measured in 15-minute intervals for 2 hours. Area under the curve was calculated by averaging the sum of all glucose per genotype. We then calculated the rate of initial drop in blood glucose after insulin administration. We limited data to the first 60 minutes after injection and modeled the exponential rate of decay in glucose for each animal as a slope. This rate was then averaged by genotype.

Twenty-four hours after fasted ITT, we collected two non-fasted blood samples: at ZT1 and ZT13. Dams were lightly anesthetized via inhaled isoflurane in a drop jar and whole blood was collected by retro-orbital bleed in a heparinized capillary tube. Blood was allowed to clot on ice for 20 minutes then was spun down in a cold centrifuge (4°C, Eppendorf microcentrifuge, model 5415R) for 20 minutes at 2000 g. Serum was pipetted off after centrifugation and stored at -80°C until further analysis.

### Serum GDF15 Quantification

GDF15 levels were quantified using maternal serum collected after ITT on E16.5 in the GDF15 and maternal comparator C57BL/6J studies. GDF15 levels were determined via ELISA according to manufacturer guidelines (R&D system, catalog # MGD150).

### Offspring Assessments

Latency to copulatory plug was defined as the number of days between the introduction of the male and appearance of a copulatory plug. Gestational age was determined as the difference between birth dates and dates of appearance of copulatory plug. Litter sizes were counted, and body weights were recorded for each pup within 24 hours of birth, PND 0.5. At PND 3.5, litter sizes were culled to 2 male and 2 female pups, to standardize the amount of nutrition/milk produced by dams and provided to pups. Survival of pups to PND 3.5 was assessed by comparing the number of pups present at PND 3.5, prior to culling, to the number present on PND 0.5 and is expressed as a percentage. Body weight was assessed for each pup on PND 0.5, 3.5, 7.5, 10.5, and 14.5. Pups were euthanized by decapitation two hours before milk collection began (PND 14.5-17.5).

### Weigh-suckle-weigh, milk volume production

On PND 10.5, we assessed milk volume production by the weigh-suckle-weigh method (30,31). Dams were weighed using an analytical balance to the nearest 10 mg and placed in a clean cage with *ad libitum* access to food and water. Pups were then weighed in aggregate and placed in a clean cage on top of a heating pad without access to food or water. Dams and pups remained separated for two hours. After two hours, weight measurements were repeated, and pups were then reintroduced to the dam’s cage. Pups remained in the dam’s cage for one hour and were allowed to nurse undisturbed. After one hour, the final weights were taken for both dams and pups in aggregate. The volume of milk produced is expressed the by average weight lost by each dam after one hour of nursing divided by the number of pups in the litter.

### Milk collection

Milk collection took place on PND 14.5-17.5. Pups were separated from dams and sacrificed two hours before milk collection began. Dams were allowed *ad libitum* access to food and water in a clean cage during that time. Dams were anesthetized with intramuscular injection of Ketamine/Xylazine (0.13g/kg body weight) into the forelimb. Once the dam was fully anesthetized, an oxytocin injection (2U per dam) was given in the forelimb muscle to promote milk ejection. Milk was collected with a pipette after manually expressing milk from nipples and stored in a 1.5 mL Eppendorf tube. Following milk collection, dams were immediately euthanized via isoflurane inhalation and secondarily with cervical dislocation.

### Milk fat percentage determinations

Whole milk was collected from dams at PND 14.5-17.5 and was stored at -80° C until analyzed. Whole milk was thawed on wet ice then homogenized by pipetting. Milk was then diluted in PBS+EDTA at a 1:3 ratio and mixed thoroughly. Capillary tubes were filled with the diluted milk solution and one end was double-sealed with crit-o-seal. Sample tubes were spun in 8 consecutive 120-second cycles in a mini hematocrit spinner (Iris Sample Processing, StatSpin CritSpin M961-122). After 16 total minutes of spinning, total fat and aqueous layers were visible within the capillary. These layers were measured using a 150mm dial caliper (General Tools, 6” Dial Caliper). Percentage of milk fat was determined based on total volume of diluted milk sample. Milk samples were analyzed in duplicate, or triplicate if milk fat percentage differed by more than 25% in the first two samples.

### Statistical Analyses

Data were analyzed in R Studio version 4.2.0 (32) and are presented as mean ± standard error (SE). Longitudinal analyses, such as food intake, body composition, and insulin tolerance testing were assessed using linear mixed effect modeling with R package lme4 (33) with random slopes and intercepts for the dam and pup with respect to time and fixed effects of genotype, age, and sex. Models for offspring body weight were assessed for interaction of sex with time and genotype but neither were significant, so sex remained a fixed effect. Pairwise values were assessed for normality by the Shapiro-Wilk test and equivalence of variance by Levene’s test. Variables that were not normally distributed or of equivalent variance underwent non-parametric testing via Mann-Whitney U test. Those that were normally distributed and of equivalent variance were assessed via Student’s *t*-test as noted in the figure legends. For this study, p-values <0.05 were considered statistically significant.

## Results

### GDF15 is elevated during pregnancy in mice

Previous work has shown that pregnancy in mice results in maternal insulin resistance (34,35). We sought to understand if GDF15 levels related to either pregnancy or a model of excess insulin resistance in pregnancy. We compared age-matched pregnant and non-pregnant females using an IP ITT on day 16.5 of pregnancy (**Figure 2A**). Consistent with prior work, we found that pregnant dams tended to be less responsive to insulin than non-pregnant females, though this did not reach statistical significance (**Figure 2A**, p=0.23 via mixed linear models). Inconsistent with Musial and colleagues, there were no significant differences in their fasting blood glucose (**Figure 2B**, p=0.20). We found that GDF15 is 49% (54 ±18.8 pg/dL) elevated in pregnant animals compared to non-pregnant mice (**Figure 2C**, p=0.007). As expected, body weights in pregnant females were 1.57± 0.55 grams heavier than non-pregnant females (**Supplemental Figure 2A**, p=0.0039).

**Figure 2:**
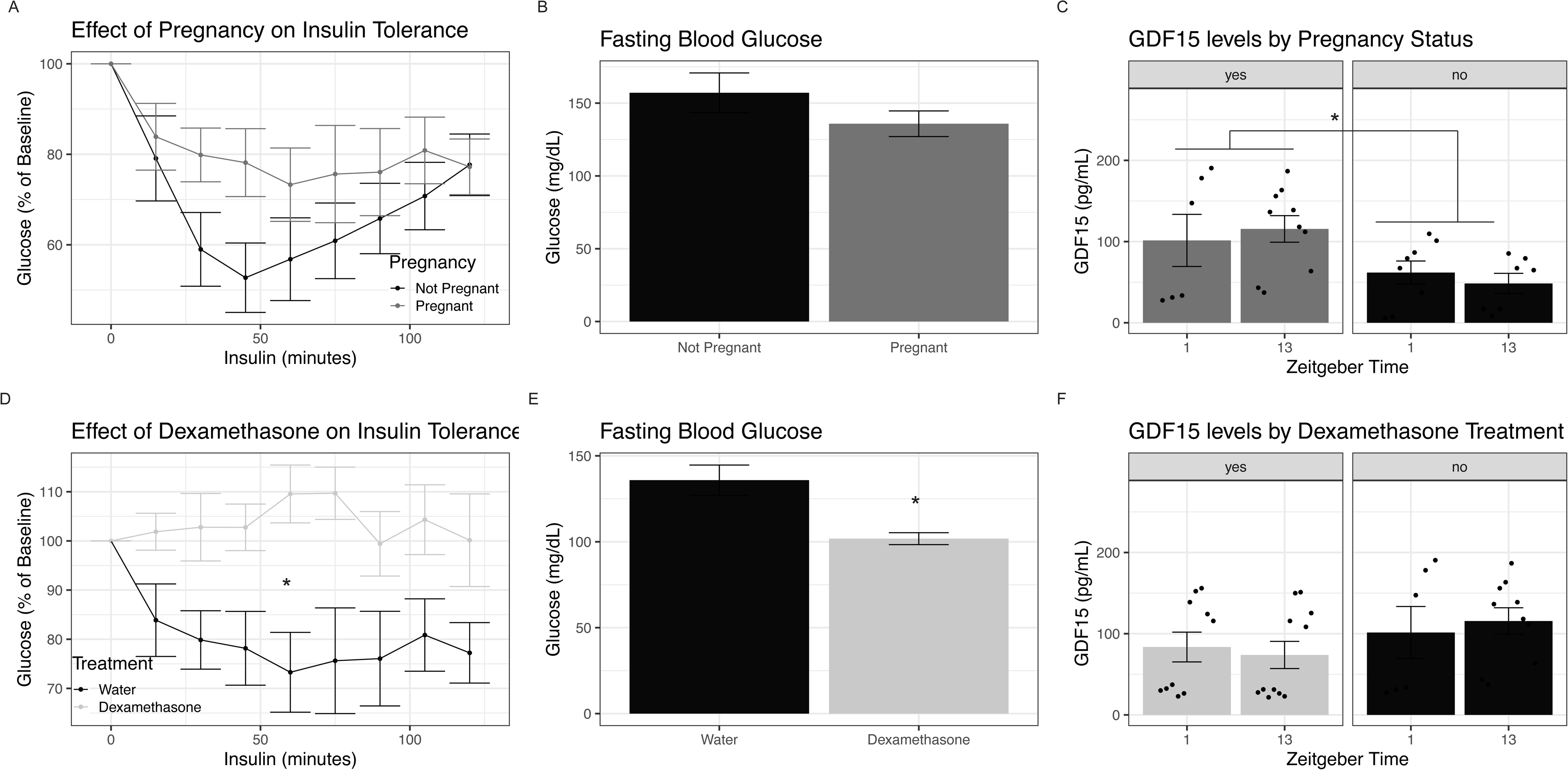
Insulin Resistance of Pregnancy Co-occurs with Elevations in GDF15. A) Intraperitoneal insulin tolerance testing on E16.5 in pregnant C57BL/6J mice given plain water and age-matched non-pregnant females. Values are relative to fasting blood glucose and were assessed using a linear mixed effect model. B) Fasting blood glucose values in pregnant dams given water and non-pregnant females, assessed using student’s T test. C) GDF15 levels at ZT1 in pregnant and non-pregnant females, assessed as paired t tests. D) Intraperitoneal insulin tolerance testing on E16.5 in pregnant dams given water or 1mg/kg dexamethasone in drinking water, assessed via linear mixed effect modeling. Values are relative to fasting blood glucose levels. E) Fasting blood glucose values in pregnant dams given plain drinking water or dexamethasone in drinking water, assessed via student’s t test. F) GDF15 ELISA evaluating serum levels at ZT1 and ZT13 in pregnant dams given plain drinking water, pregnant dams given dexamethasone in drinking water, assessed as paired t tests. * indicates p<0.05

To enhance insulin resistance in pregnancy, we leveraged prior work from our lab which has demonstrated that administering the glucocorticoid dexamethasone (dex) in the drinking water impairs insulin sensitivity in non-pregnant mice (36,37). We treated dams with 1 mg/kg dexamethasone one week before mating and throughout pregnancy. We compared dexamethasone-treated dams to age-matched pregnant dams who were provided untreated drinking water. We found that dexamethasone dams did not respond to insulin compared to pregnant dams with plain drinking water (**Figure 2D**, p_dex*time_=0.02 via linear mixed effect models). Dexamethasone-treated dams had 33% lower fasting blood glucose (**Figure 2E**, p_dex_=0.007) consistent with our findings in non-pregnant mice. GDF15 levels were not further increased by dexamethasone administration in pregnant dams (**Figure 2F**, p=0.11). Body weights in pregnant dams were 2.77±0.58 grams less in those treated with dex compared to untreated dams (**Supplementary Figure 2B**, p<0.0001). Based on these data we conclude that while GDF15 is related to pregnancy, it is not elevated in insulin resistant dexamethasone treated dams.

### Gdf15^-/-^ dams have normal weight gain and modestly reduced food intake during pregnancy and lactation

To evaluate the role of *Gdf15* ablation in maternal food intake and body weight accretion during pregnancy, we mated *Gdf15^+/+^* dams with *Gdf15^+/+^* males and compared them to *Gdf15^-/-^* mated pairs (**Figure 1B**). Dam body weight and food intake were measured weekly, beginning one week before mating and continued until pups reached 14 days of age (PND14.5).

*Gdf15^-/-^* dams consumed similar cumulative kilocalories during the prenatal period (**Figure 3A**, p=0.52). They also had a similar weight change when compared to *Gdf15^+/+^*dams during the course of pregnancy (**Figure 3B**, p=0.99). Both genotypes consumed similar calories weekly (**Figure 3E**, p_genotype_=0.23). Both genotypes had a rapid increase in food intake in the final week of pregnancy, with smaller increases in the *Gdf15^-/-^*dams. In the postnatal period, cumulative food intake was similar between genotypes (**Figure 3C**, p=0.94). *Gdf15^-/-^* dams had 54% lower postnatal weight loss than *Gdf15^+/+^* dams with high levels of variability, but this failed to reach statistical significance (**Figure 3D**, p=0.20; **Figure 3F**). This suggests that *Gdf15* is not a major determinant of either body weight or food intake during the first pregnancy in mice.

**Figure 3:**
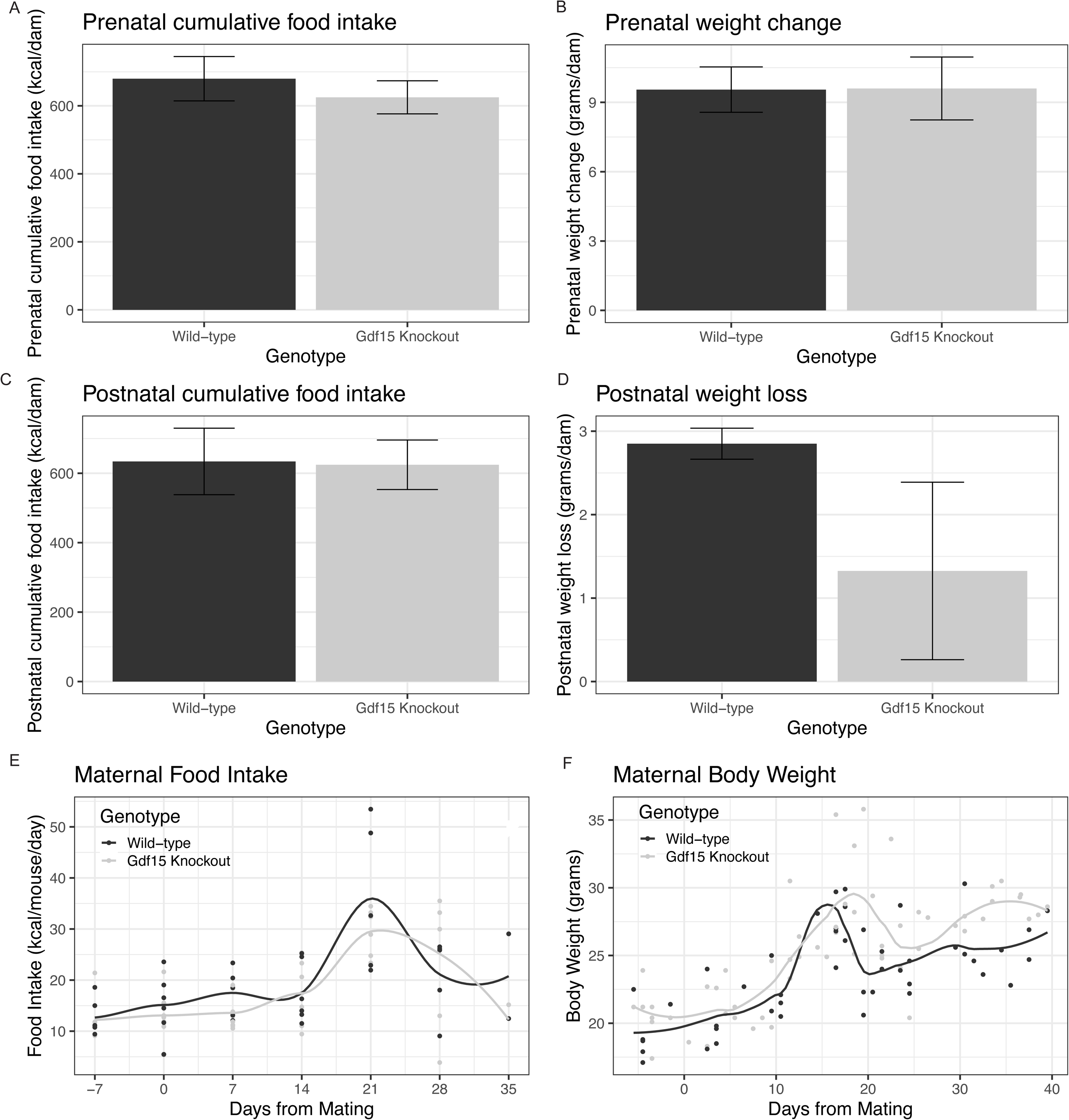
Gdf15 Knockout Does Not Impact Food Intake or Body Weight During Mouse Pregnancy. A) Cumulative food intake during the prenatal period (pre-mating through final measurement before birth), assessed via Student’s *t* test. B) Weight gained during prenatal period, assessed via student’s t test. C)Postnatal cumulative food intake (after birth of pups-end of experiment), assessed via Student’s *t* test. D) Weight lost in the postnatal period, assessed via Students’ *t* test. E) Plot of the weekly food intake in both genotypes from 1 week before mating until end of the experiment. F) Plot of maternal body weight throughout the experimental period.

### Gdf15^-/-^ dams have normal insulin tolerance during pregnancy

On Gestational day 16.5, we conducted an intraperitoneal insulin tolerance test to assess the effect of *Gdf15* ablation on maternal insulin sensitivity during pregnancy (**Figure 4A**). Fasting blood glucose was slightly but insignificantly lower in *Gdf15^-/-^* dams compared to *Gdf15^+/+^* dams (**Figure 4B**, p = 0.20). Overall, linear mixed effect modeling revealed no effect of genotype (p_genotype_ = 0.71). This was confirmed by determining the area under the curve, again showing similar responses (**Figure 4C**, p=0.74). Often an informative measure of the insulin response is the initial rate of drop of blood glucose. The initial rate of glucose decline was 9.3% less in *Gdf15^-/-^* dams compared to *Gdf15^+/+^* dams, however, this did not reach statistical significance (**Figure 4D**, p=0.082). These data suggest that ablation of *Gdf15* is not sufficient to substantially alter insulin sensitivity in the pregnant mouse.

**Figure 4:**
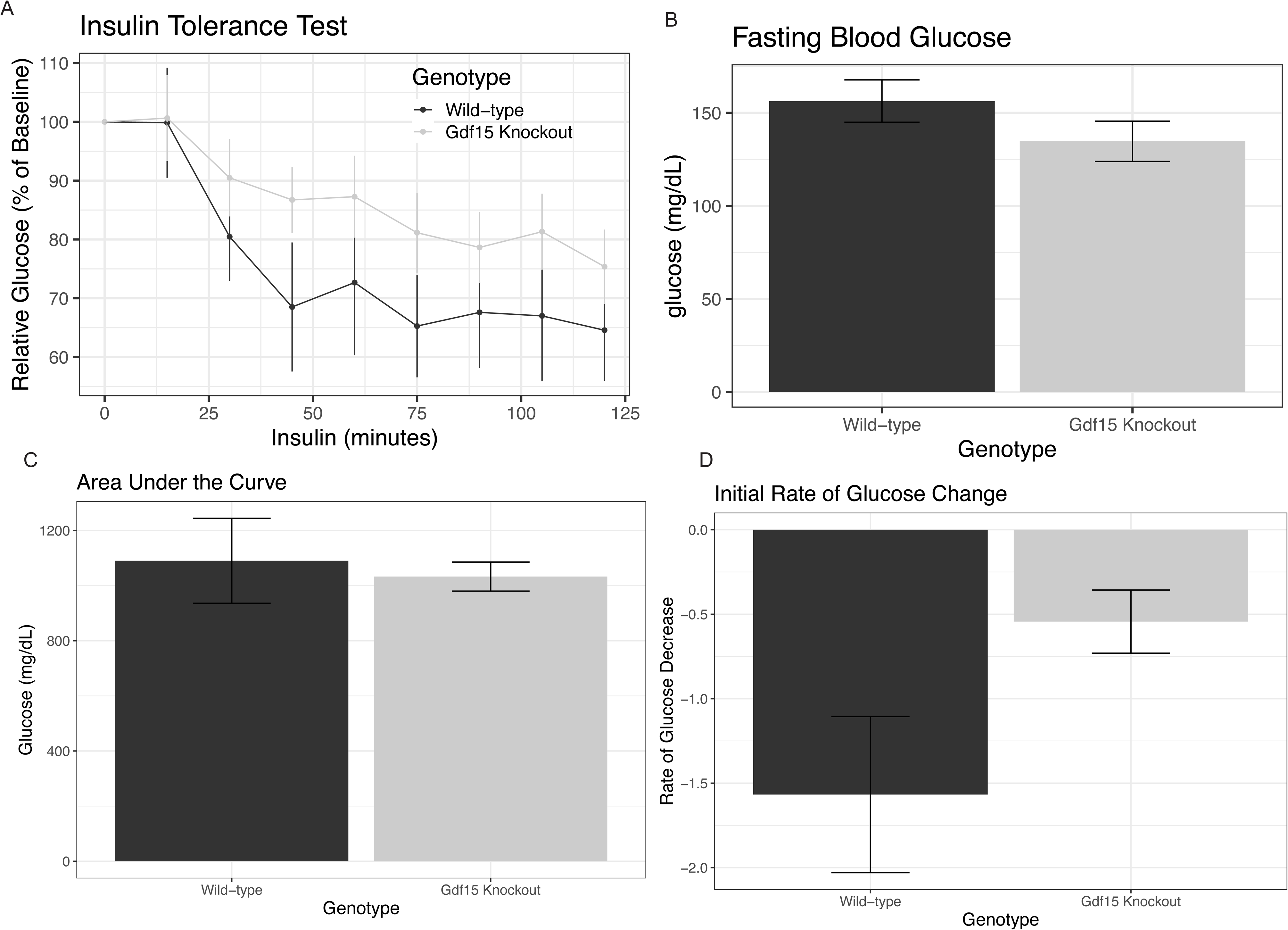
Gdf15 Knockout Has No Effect on Gestational Insulin Tolerance. A) Intraperitoneal insulin tolerance test in *Gdf15^+/+^* and *Gdf15^-/-^*dams at E16.5. Values are relative to fasting blood glucose levels. Assessed via linear mixed effects modeling. B) Fasting Blood glucose levels in dams, assessed by students t test. C) Area under the curve defined as sum of all glucose values for each animal, assessed by Student’s *t* test. D) Rate of drop in blood glucose in the first hour of the insulin tolerance test, assessed by Student’s *t* test.

### Gdf15^-/-^ dams have normal fertility, gestational age, post-natal survival, and pup birth weights

To understand the role of *Gdf15* knockout on gestational health, we calculated latency to plug, gestational age, and litter size. To assess early postnatal outcomes in the pups we evaluated birth weight and 3-day survival. The latency to copulatory plug was similar between genotypes, averaging 3 days (**Figure 5A**, p=0.74). Gestational age at delivery was similar between genotypes, averaging 20 days (**Figure 5B**, p=0.76). Pups from *Gdf15^-/-^* dams were 3.4% smaller than those from *Gdf15^-/-^* dams (**Figure 5C**, p=0.05). The total number of pups born per litter was 27% greater in *Gdf15^-/-^* dams (1.6 pups greater on average) compared to *Gdf1^+/+^* dams (**Figure 5D**, p=0.15). When comparing litter size, counting only pups alive at birth, that difference was reduced to 7.8% larger (**Figure 5E**, p=0.70, or 0.46 pups/litter greater on average). The total pups who were born alive that lived to PND 3.5 was variable within genotypes, resulting in 91.7% survival for *Gdf15^+/+^* dams and 90% for *Gdf15^-/-^* dams which was not statistically significant (**Figure 5F**, p=0.99). Together these data show that aside from modest decreases in birthweights, *Gdf15^-/-^*mice are similarly fertile, and carry pregnancies to a similar effectiveness as their wild-type counterparts.

**Figure 5:**
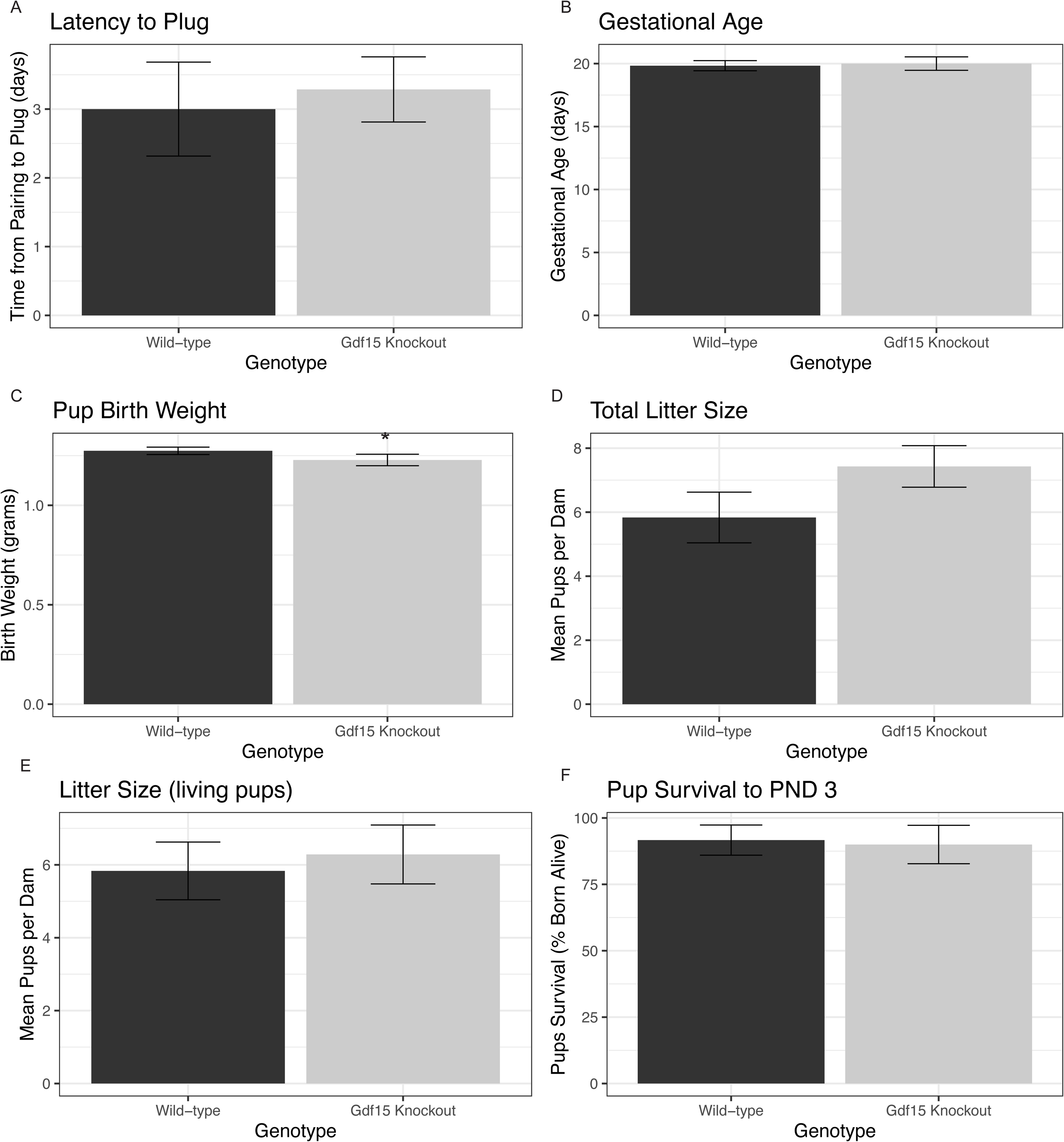
Offspring Birth Weight is Reduced in Gdf15 Knockout Pregnancies. A) Latency to copulatory plug (time from introduction of male into cage until copulatory plug is discovered), assessed via student’s t test. B) Gestational age in days, calculated as the number of days from appearance of copulatory plug until birth of the litter. Assessed via Mann-Whitney test. C) Average birth weight of pups, calculated as the average birth weight for each dam, then averaged by genotype. Assessed by Student’s *t* test. D) Total litter size (including those born dead), assessed via student’s t test. E) Number of live pups born per litter, assessed via student’s t test. F) Percentage of pups in each litter who were dead by postnatal day 3.5, assessed by Mann Whitney test. * indicates p<0.05

### Gdf15^-/-^ dams have no differences in milk production or milkfat percentage

To determine the effect of *Gdf15* knockout during pregnancy on lactation, we assessed milk volume at PND 10.5 as described by Boston and colleagues (30). We found no differences between *Gdf15^+/+^and Gdf15^-/-^* dams in the volume of milk produced. The amount of weight lost by dams after nursing (**Figure 6A**, p=0.7) and weight gained by pups during nursing (**Figure 6B**, p=0.7) was similar between genotypes, though highly variable between dams. Next, we evaluated whether the major macronutrient in milk, fat, was changed by *Gdf15* knockout. To do this, we collected whole milk between PND 14.5-17.5 and evaluated milk fat percentage. We found that milk fat percentage was similar between genotypes (**Figure 6C**, p=0.93). Despite reduced maternal levels of GDF15 in the *Gdf15^-/-^* dams during pregnancy, gestational mammary gland development, and lactation there is no apparent impact on milk volume or milk fat content.

**Figure 6:**
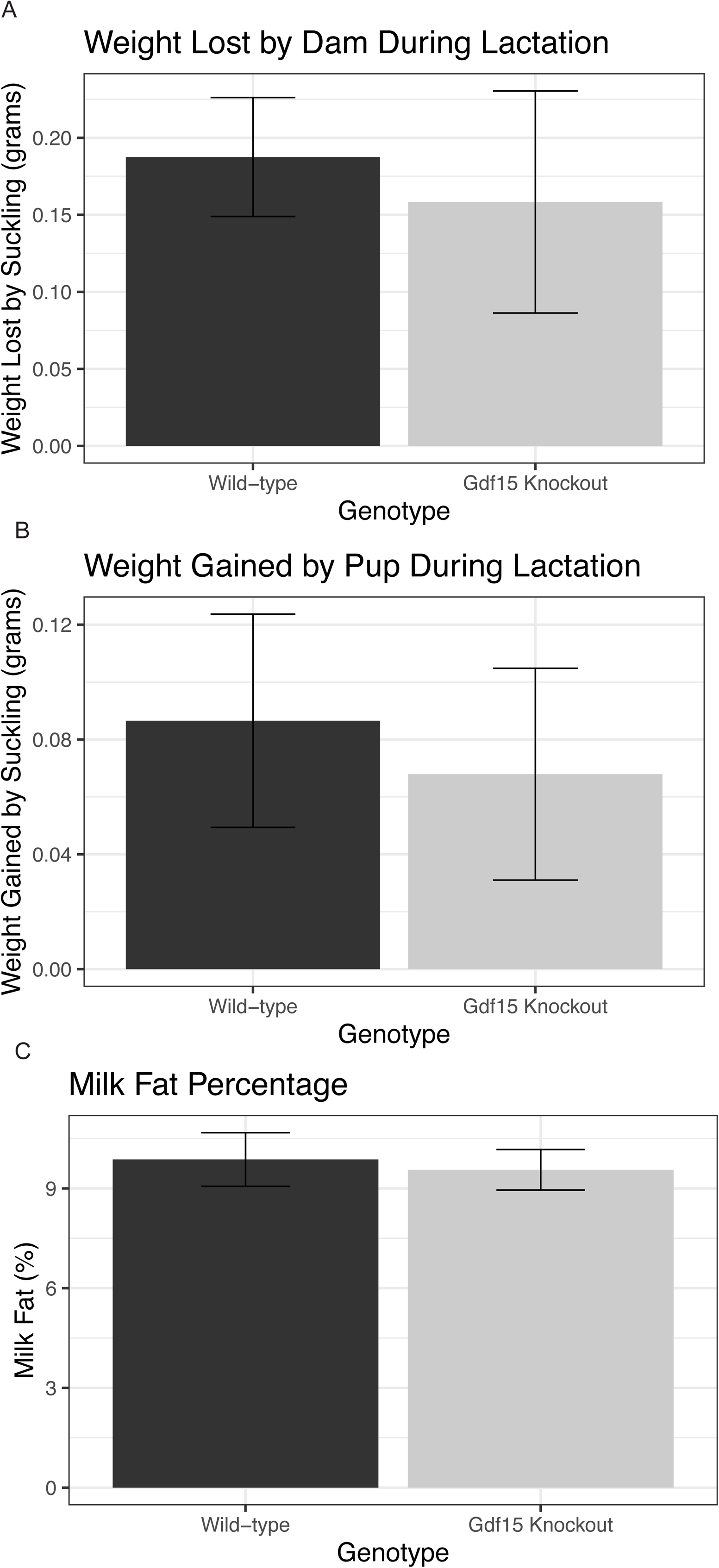
Milk volume and Milkfat Percentage Are not Changed in Gdf15 Knockout Dams. A) Total mass (in grams) lost by dam during the suckling period of the weigh-suckle-weigh test on PND10.5, assessed by student’s t test. B) Total mass (in grams) gained cumulatively between all pups in the litter during suckling period during weigh-suckle-weigh test, assessed by Mann Whitney test. C) Percentage of fat found in mouse milk collected PND 14-16.5, assessed by student’s t test.

### Gdf15^-/-^ pups accrete body mass at similar rates compared to Gdf15^+/+^ pups

To assess the effect of *Gdf15* knockout during pregnancy and lactation on early pup postnatal growth, we weighed male and female offspring of *Gdf15^+/+^* and *Gdf15^-/-^* dams on PND 0.5, 3.5, 7.5, and 14.5. We used linear mixed effect modeling which detected no differences in body weight between birth and 14 days of age in *Gdf15^+/+^* and *Gdf15^-/-^* pups (**Figure 7A**, p_genotype_=0.81 after adjusting for sex differences). There was also no statistically significant modifying effect of sex on body weight from birth to PND 14.5 (p_sex_=0.16). Therefore, consistent with similar milk production and composition, we did not detect any effects of *Gdf15* ablation on perinatal growth.

**Figure 7:**
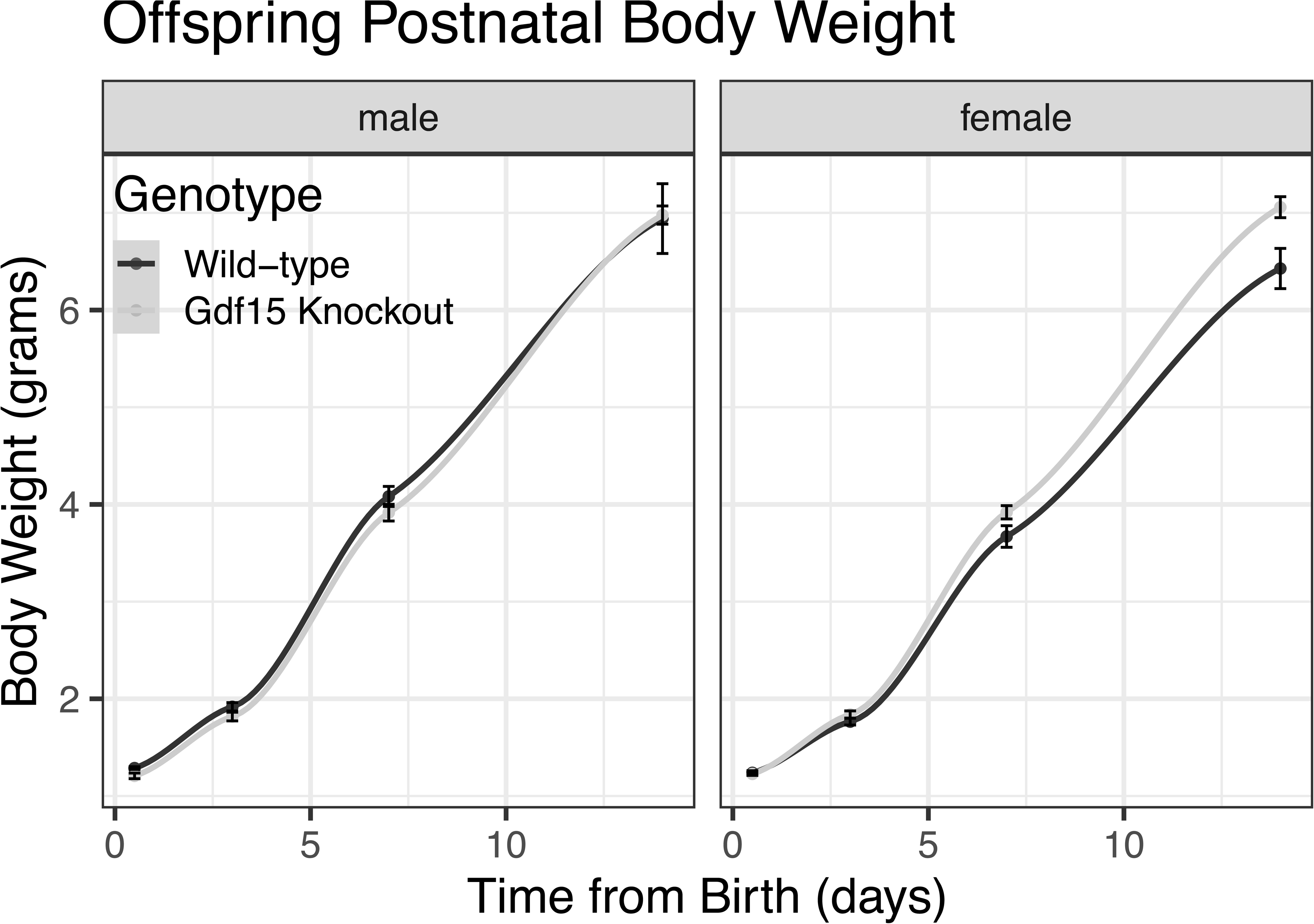
Offspring Postnatal Growth is Normal in Gdf15 Knockout Litters. A) Postnatal bodyweight measurements from birth through PND14.5 in male and female pups, assessed via linear mixed effect models.

## Discussion

GDF15 has recently been linked to several complications of pregnancy in addition to its role in signaling somatic stress throughout the body. Pregnancy itself is an oft-underappreciated stressor on the body, an effect that is consistent with elevations in GDF15. The goal of this study was to understand the role of GDF15 in gestational health. To date, there are very few studies that evaluate GDF15 in human pregnancy. Circulating GDF15 levels during pregnancy is associated with body weight and BMI is inconsistent in human cohorts. Elevated circulating levels of GDF15 have been documented in expectant parents with normal weight status compared to those with obesity (7). GDF15 has also been negatively associated with total gestational weight gain (22). Because of these associations in human pregnancy studies with lower body weights and gestational weight gain, we anticipated unrestrained food intake and weight gain in our murine model. However, the lack of prominent changes in gestational outcomes, is nonetheless novel in GDF15 literature. Previous reports of *Gdf15* or *Gfral* null studies have generally not reported pregnancy or gestational outcomes during breeding or maintenance, but only describe differences as adults when used in experimental models. To our knowledge, the only study noting major changes in gestational outcomes assessed transgenic expression of human *GDF15* in mice and found that there was early mammary gland involution, reduced milk production, reduced survival in pups, and lower weight gain in the postnatal period in offspring born to transgenic dams (38). The unexpected lack of evidence that loss of GDF15 is related to pregnancy related eating and weight gain may also be due to a lower level of residue homology between human and mouse *Gdf15* (39). It could also explain why using transgenic, human *GDF15* was associated with pregnancy related body weight and lactational differences when we saw none (38). Previous work shows that external administration of GDF15, similar to the rising levels accompanying pregnancy, in mice results in reductions in food intake (15,40). Our current study found that ablation of *Gdf15* and the resulting loss of GDF15 in maternal circulation (**Supplementary Figure 1A**) does not result in any differences in body weight accretion during the prenatal period and resulted in non-statistically significant higher body weights during the postnatal period in mice, with only small reductions in pup birth weight. This suggests that GDF15 in pregnant mice is altered, but it is not necessary for changes in weight accretion during a normal mouse pregnancy. It is possible that under conditions of elevated somatic stress other than pregnancy, GDF15 plays a larger role. It is also possible that the deletion of GDF15 during mouse pregnancy is of little effect because GDF15 in circulation increases only modestly in rodents (2-fold) compared to humans and primates (75-100 fold in primates and humans, respectively) (41).

Taken together, the lack of evidence of differences in food intake, body weight, insulin sensitivity, and lactation in our *Gdf15* null model suggests that there may be a threshold effect for GDF15 during pregnancy. Only those studies that overexpress, deliver exogenous, or induce long-term highly disruptive stressors to their model show differences in food intake and body weight related to GDF15 levels. Therefore, it might be that typical pregnancy-related inductions of GDF15 in rodents are insufficient to meet the threshold to elicit an effect. *Gdf15* may act as a less acute stressor during pregnancy and more as a long-term indicator of feto-placental implantation. It could also imply that in observational human studies, GDF15 is a biomarker of pregnancy related complications but not part of a causal pathway.

There are several limitations to our study. Murine pregnancy is not entirely comparable to human pregnancy. The majority of human pregnancies are singleton and mice are multi-parous. Murine placental structure is also different when compared with human placentae, in the level of invasion of the tissue into the maternal uterus and the structure of the zones of the placenta itself (42). The approach we took eliminated feto-placental contribution of GDF15 to maternal serum during pregnancy using homozygous breeding pairs. As a result, all knockout pups had knockout dams and sires, and all wild-type pups had wild-type dams and sires. In the offspring from these like-genotype pairings, we did not detect any differences in growth. A larger sample size could have provided more statistical power to detect differences in the outcomes evaluated. It is also possible that compensatory adaptations, such as other feeding hormones occur alongside with life-long loss of GDF15. We also followed the pups for a relatively short period of time after birth (PND 14.5). Any effect that would have manifested after the second week of life was not evaluated. Finally, we did not evaluate two other GDF15-associated complications, hypertension, or nausea-related behavior in these mice.

In contrast to the human findings, this study had several strengths including strong environmental, genetic, and experimental consistency. Dams and sires were homozygous, they were derived from heterozygous crosses to limit genetic drift. In contrast to human observational studies demonstrating connections to pregnancy complications, we do not observe detect differences in litter sizes, glucose homeostasis, or gestational weight gain in the knockout mice. Ours is the first report of the loss of GDF15 on pregnancy outcomes in a preclinical model and provides strong evidence that GDF15 levels are not critical for pregnancy related body weight gain, increases in food intake, or early postnatal offspring health.

## Conclusion

Despite the well-known rise in GDF15 during mouse and human pregnancy, we found no evidence that *Gdf15* ablation during mouse pregnancy and lactation causes metabolic, body weight, appetite, or lactational differences compared to age-matched *Gdf15^+/+^* dams. In the neonatal period, we did not observe any differences in survival, gestational age, litter size and only modest birth weight reductions between genotypes. Despite monitoring growth for two weeks after birth, body weight accretion in *Gdf15^-/-^* pups of either sex was indistinguishable from age-matched *Gdf15^+/+^* pups. Further studies with larger sample sizes are needed to confirm these findings.

## Disclosures

None

## Author contribution

MCM, NEH , BEG, and DB conceptualized experiments. Data were curated by MCM, NEH, JRR, and HS. MCM completed the analysis and wrote the original draft of the manuscript. All authors contributed to the review and editing of the manuscript and approved its final format before submission.

## Funding

MCM is supported by a Rackham Merit Fellowship. This study was supported by funds from the University of Michigan (MCubed to DB) and the National Institutes of Health (DK107535 and a Pilot and Feasibility Grant from P30DK020572 to DB, and DK K08102526 to BG). This study was supported by core services provided by the Michigan Nutrition and Obesity Research Center (DK089503).

## Notes

https://github.com/BridgesLab/Developmental-Obesity

### Competing Interest Statement

The authors have declared no competing interest.

