## Supplementary figures and images for "GDF15 knockout does not substantially impact perinatal body weight or neonatal outcomes in mice"

### Supplementary Figures

Supplementary Figure 1:

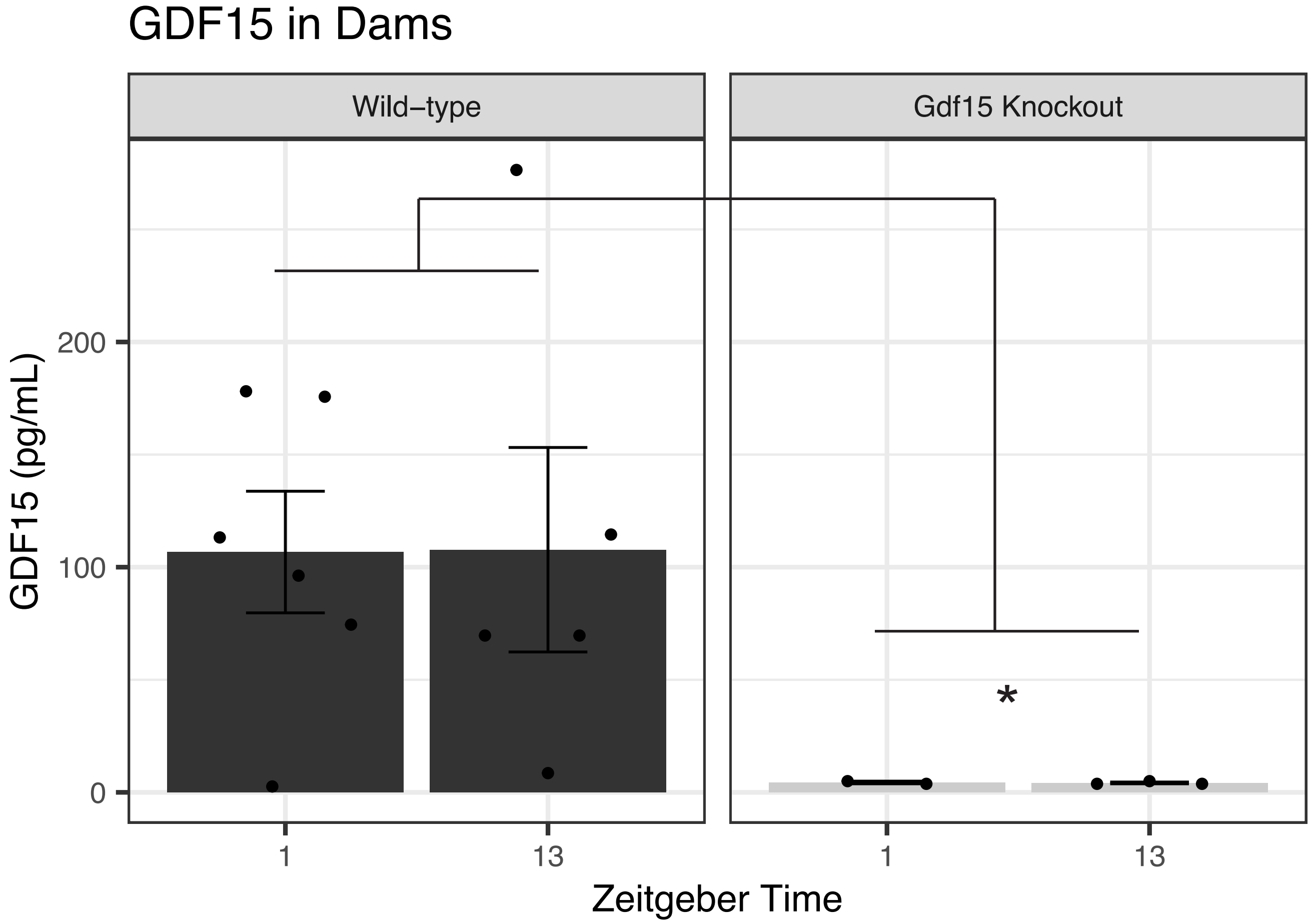

Supplementary Figure 2:

A

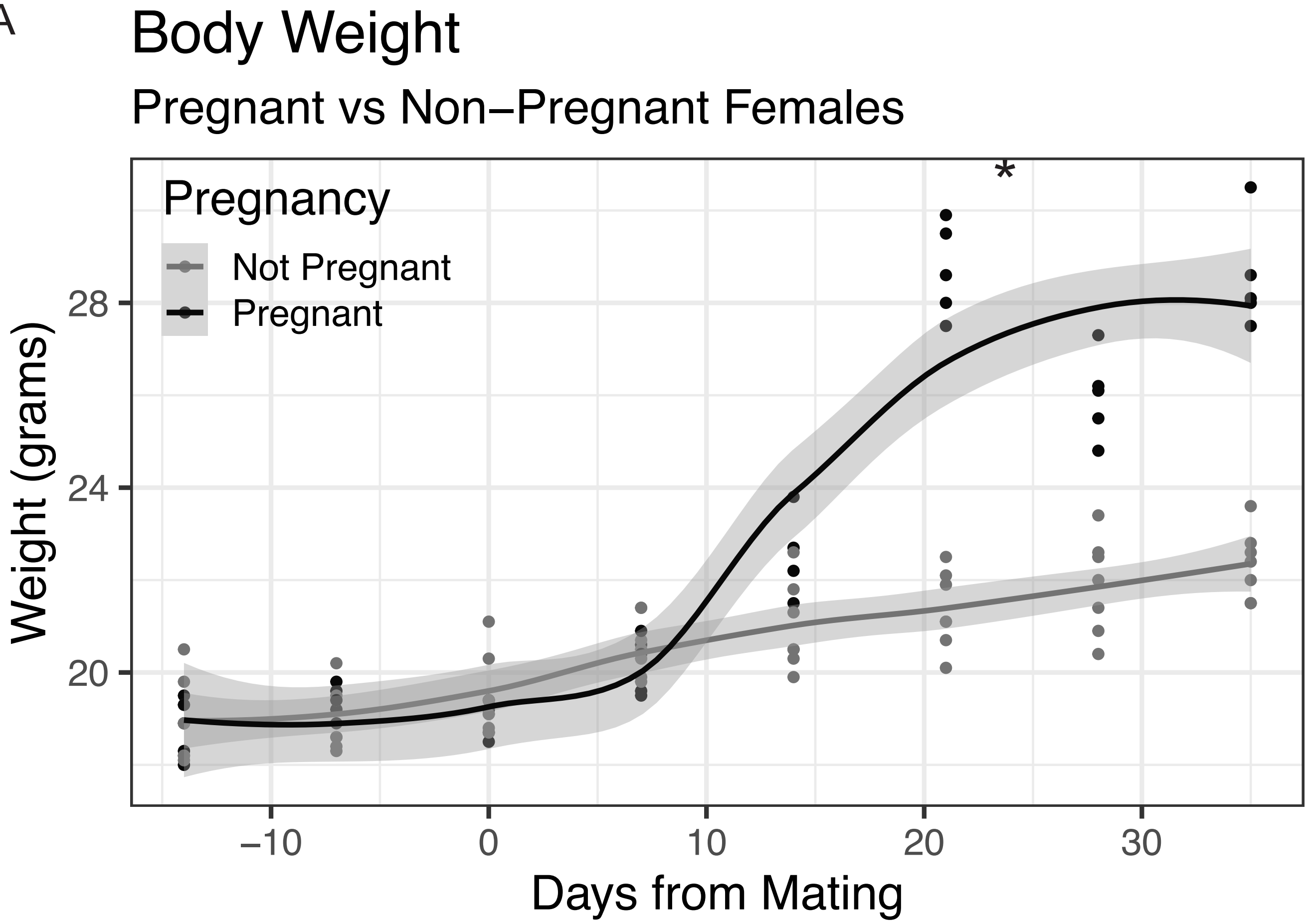


B

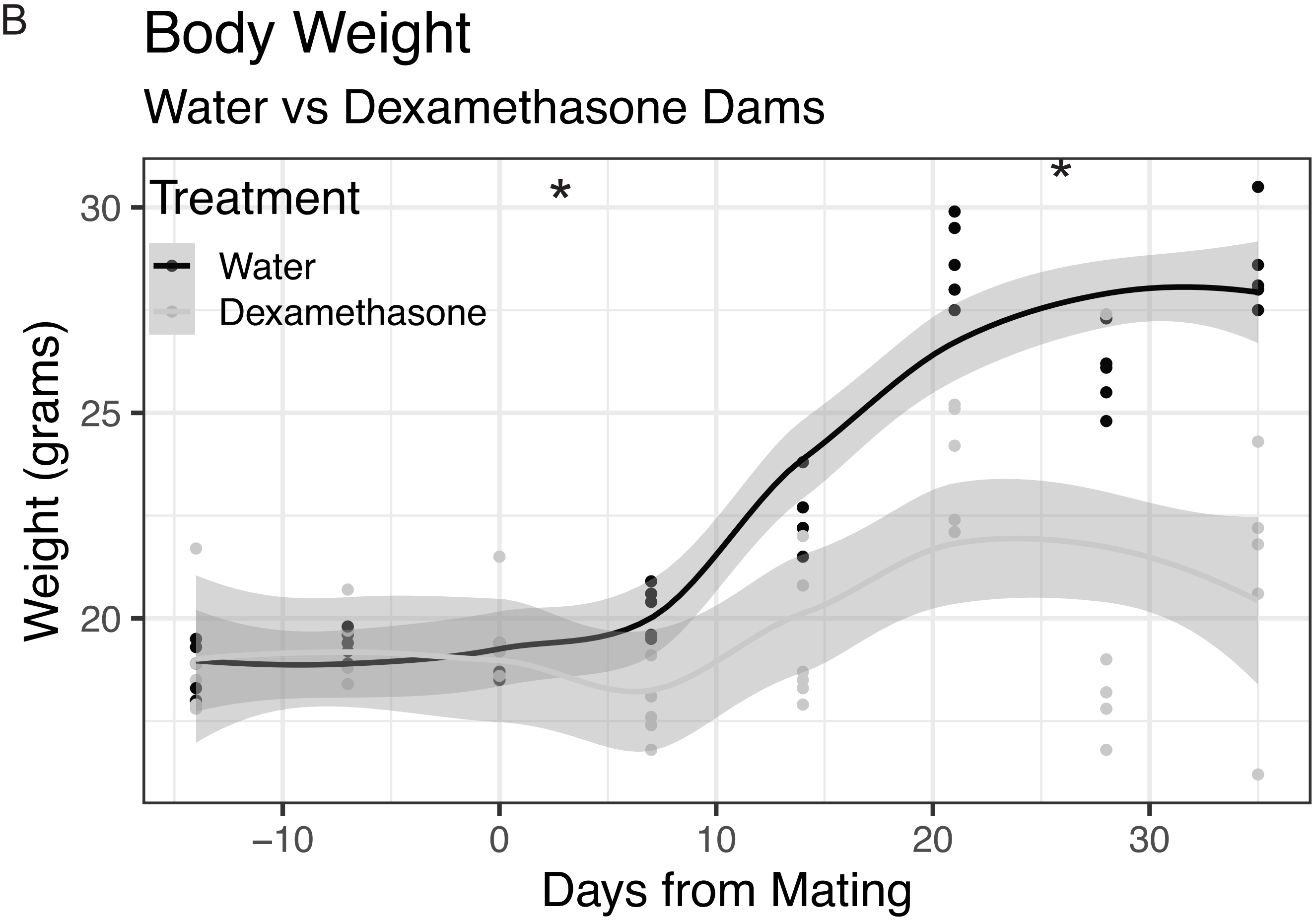
